# Naturalistic coding of working memory in primate prefrontal cortex

**DOI:** 10.1101/2020.06.19.162446

**Authors:** Megan Roussy, Rogelio Luna, Lyndon Duong, Benjamin Corrigan, Roberto A. Gulli, Ramon Nogueira, Ruben Moreno-Bote, Adam J. Sachs, Lena Palaniyappan, Julio C. Martinez-Trujillo

**Affiliations:** Department of Physiology and Pharmacology, the University of Western Ontario, London, ON, Canada; Robarts Research Institute, the University of Western Ontario, London, ON, Canada; Zuckerman Mind Brain Behaviour Institute, Columbia University, New York, NY, USA; Center for Theoretical Neuroscience, Columbia University, New York, NY, USA; Center for Neural Science, New York University, New York, NY, USA; Center for Brain and Cognition, and Department of Information and Communication Technologies, Universitat Pompeu Fabra, Barcelona, Spain; Serra Húnter Fellow Programme, Universitat Pompeu Fabra, Barcelona, Spain; The Ottawa Hospital, University of Ottawa, Ottawa, ON, Canada; Department of Psychiatry, the University of Western Ontario, London, ON, Canada; Lawson Health Research Institute, London, ON, Canada; Brain and Mind Institute, the University of Western Ontario, London, ON, Canada

## Abstract

The primate lateral prefrontal cortex (LPFC) is considered fundamental for temporarily maintaining and manipulating mental representations that serve behavior, a cognitive function known as working memory^1^. Studies in non-human primates have shown that LPFC lesions impair working memory^2^ and that LPFC neuronal activity encodes working memory representations^3^. However, such studies have used simple displays and constrained gaze while subjects held information in working memory^3^, which put into question their ethological validity^4,5^. Currently, it remains unclear whether LPFC microcircuits can support working memory function during natural behavior. We tested macaque monkeys in a working memory navigation task in a life-like virtual environment while their gaze was unconstrained. We show that LPFC neuronal populations robustly encode working memory representations in these conditions. Furthermore, low doses of the NMDA receptor antagonist, ketamine, impaired working memory performance while sparing perceptual and motor skills. Ketamine decreased the firing of narrow spiking inhibitory interneurons and increased the firing of broad spiking cells reducing population decoding accuracy for remembered locations. Our results show that primate LPFC generates robust neural codes for working memory in naturalistic settings and that such codes rely upon a fine balance between the activation of excitatory and inhibitory neurons.

## Main

The lateral prefrontal cortex (LPFC, areas 8A/9/46) is a part of the granular neocortex that emerged during evolution of anthropoid primates^6^. LPFC is thought to encode short term memory representations dissociable from sensory and motor signals that serve behavioral goals, an essential cognitive function known as working memory (WM)^3,4,6,7^. Various studies in macaque monkeys have shown that the activity of LPFC neurons encode visuospatial information held in WM^3,8,9,10^. Such studies have employed behavioral tasks involving simple visual displays relative to the complexity of natural scenes and have strictly controlled for eye movements^3,4,5^. However, in real-life settings, WM representations must be held during dynamic viewing of natural scenes through saccades. The activity of LPFC neurons has been shown to encode visual stimuli^11^, raising the question of whether LPFC microcircuits can support WM function in ethologically valid settings. Here, we aimed to clarify this issue.

We used a virtual reality engine to build a virtual arena featuring a naturalistic visual scene. We trained two rhesus monkeys (Macaca mulatta) on a visuospatial WM task that took place in this arena (Fig. 1a, b). As during natural behavior, animals were permitted free visual exploration (unconstrained eye movements), as well as spatial navigation using a joystick. During task trials, a target was presented for 3 seconds at 1 of 9 locations in the arena. The target then disappeared during a 2 second delay epoch. During the target and delay epoch, navigation was disabled.

**Figure 1.**
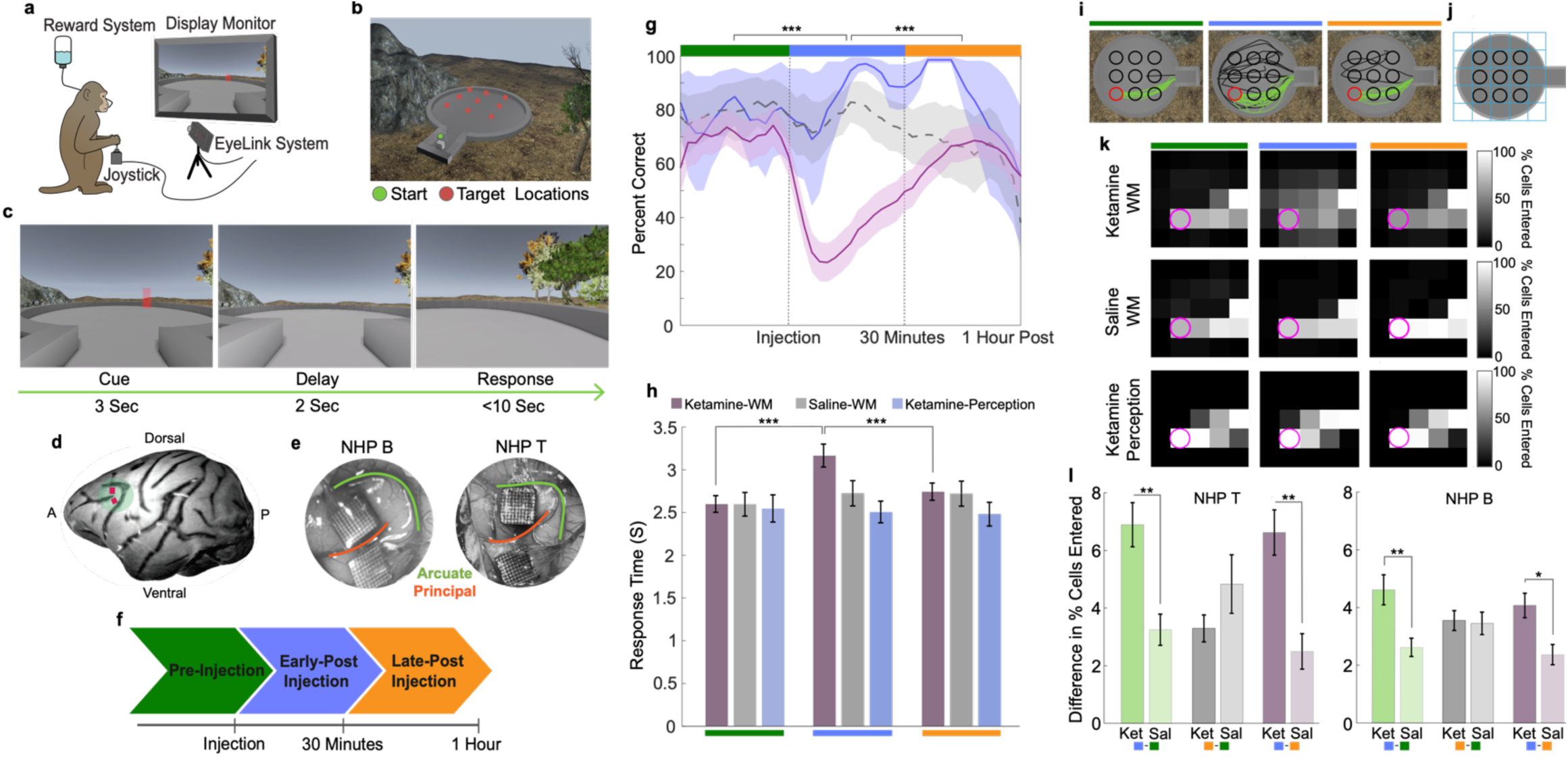
Virtual working memory task and behavioural performance. **a**, Illustration of experimental setup. **b**, Overhead view of task arena in virtual environment. **c**, Trial epoch timeline. **d**, Depiction of Utah array locations. **e**, Surgical images of Utah arrays in LPFC. **f**, Injection period timeline. Data from pre-injection period represented by green, early post-injection period by blue, and late post-injection period by orange. **g**, Average percent of correct trials for ketamine-WM sessions (pink), saline-WM sessions (grey), and ketamine-perception sessions (blue). **h**, Average response time for correct trials for all session types. Data points represent values per target location for each session. **i**, Trajectories to example target location (red) in one ketamine-WM session for correct (green) and incorrect (black) trials. **j**, Task arena divided into 5×5 grid. **k**, Percent of trials in which each cell of the arena is entered for example target location (pink) averaged over sessions. **l**, Average difference (increase) in percent of trials in which cells are entered between injection periods (green = early post-injection – pre-injection; grey = late post-injection – pre-injection; purple = early post-injection – late post-injection) compared between ketamine-WM and saline-WM sessions. All error bars are SEM. *<0.05, **<0.01, ***<0.001.

Subsequently, navigation was enabled, and animals were required to virtually approach the target location within 10 seconds to obtain a juice reward (Fig. 1c). We recorded neural activity during this task using two 96-channel microelectrode arrays (Utah Arrays) implanted in the LPFC (Fig. 1d, e)^12^.

In order to assess whether LPFC activity is causally linked to WM performance, we administered ketamine, a non-competitive N-methyl-D-aspartate receptor (NMDAR) antagonist. NMDARs are evidenced to be critically involved in balancing prefrontal circuit interactions between pyramidal cells and inhibitory interneurons that are crucial for WM processing^13,14,15,16^. Ketamine is reported to impair WM performance through primarily blocking NMDAR which are highly expressed in the human prefrontal cortex^16,17,18,19^. Antagonism of these subunits is also sufficient to perturb LPFC WM signals^14^. Accordingly, it is reasonable to assume that low doses of systemically administered ketamine would produce the greatest effect on prefrontal neural activity^20,21,22^.

We recorded neural responses during the task in three blocks of trials: before subanesthetic ketamine (0.25 mg/kg-0.8 mg/kg) or saline injection (pre-injection period), 30 minutes post injection (early post-injection period), and up to 1-hour post injection (late post-injection period) (Fig. 1f). In some sessions, we use a control task in which targets remain onscreen for the duration of the trial (ketamine-perception variant). Here, the animals did not have to remember the target location; therefore, WM was not required to complete the trials. This control variant of the task allows us to separate the effect of ketamine on WM function from potential effects on processes like perception and movement.

### Behavioral performance

Both animals performed significantly above chance (∼11%, nine locations) on all task variants before ketamine injections (pre-injection period, *p*<0.001) indicating proficiency in the task. In ketamine-WM sessions, performance decreased significantly during the early post-injection period compared to the pre-injection period (ANOVA, post hoc, *p*<0.0001), to subsequently recover during the late post-injection period compared to the early post-injection period (*p*<0.0001). Performance did not significantly change between injection periods in saline-WM sessions (ANOVA, *p*=0.075). Importantly, ketamine injections did not significantly alter performance between injection periods in perception sessions (ANOVA, *p*=0.786) indicating that the ketamine-induced performance deficit was specific to the WM task (Fig. 1g) (see data per non-human primate (NHP) in Extended Data Fig. 1 a, b). Navigation time to the remembered target location increased significantly after ketamine injection compared to the pre-injection period (ANOVA, post hoc, *p*<0.0001) and decreased in the late post-injection period compared to the early post-injection period (*p*<0.0001). No significant changes were found between injection periods in saline-WM (ANOVA, *p*=0.186) or ketamine-perception sessions (ANOVA, *p*=0.800) (Fig. 1h; see data per NHP in Extended Data Fig. 1 c, d).

Trajectories to remembered targets also became more dispersed after ketamine injections (Fig. 1i). To quantify this observation, we divided the task environment into a 5×5 grid creating 25 regional cells (see Fig. 1j) and calculated the percent of trials in which each cell was entered during navigation to a target location (Fig. 1k). The difference in the percent of cells entered between pre- and post-injection periods in ketamine-WM and saline-WM sessions was then calculated. In ketamine-WM sessions, more cells were visited in the early post-injection compared to the pre-injection period relative to saline-WM sessions (post hoc, NHP T, *p*=0.002; NHP B, *p*=0.004). Fewer cells were visited in the late post-injection period compared to the early post-injection period in ketamine-WM sessions compared to saline-WM sessions (post hoc, NHP T, *p*=0.001; NHP B, *p*=0.044) (Fig. 1l). We did not observe significant dispersion of the trajectories in the ketamine-perception task. These results indicate that ketamine selectively impaired the animals’ ability to maintain the location of the target in WM.

### Tuning of single neurons

To investigate the neural correlates of the behaviors illustrated in Fig. 1, we recorded the activity of 2906 units (1814 single neurons and 1092 multiunits) during 17 ketamine-WM sessions (8 in NHP T, 9 in NHP B). We recorded an additional 1117 units (674 single units and 443 multiunits) during seven saline-WM sessions (3 in NHP T, 4 in NHP B). Single neurons exhibited spatial tuning during the delay epoch in the pre-injection period (example neurons in Fig. 2a, b). At a population level, during ketamine-WM sessions, the proportion of spatially tuned neurons significantly decreased in the early post-injection period compared to the pre-injection period (ANOVA, post hoc, *p*=0.005) and significantly increased in the late post-injection period compared to the early-post injection period (ANOVA, *p*<0.0001) (Fig. 2c; see data per NHP in Extended Data Fig. 2a, b). There were no significant differences in the proportion of tuned single neurons between injection periods during saline-WM sessions (ANOVA, *p*=0.088) (Fig. 2d). These results demonstrate that single neurons in LPFC encode spatial WM signals in naturalistic conditions and that low doses of ketamine significantly impair single neuron tuning.

**Figure 2.**
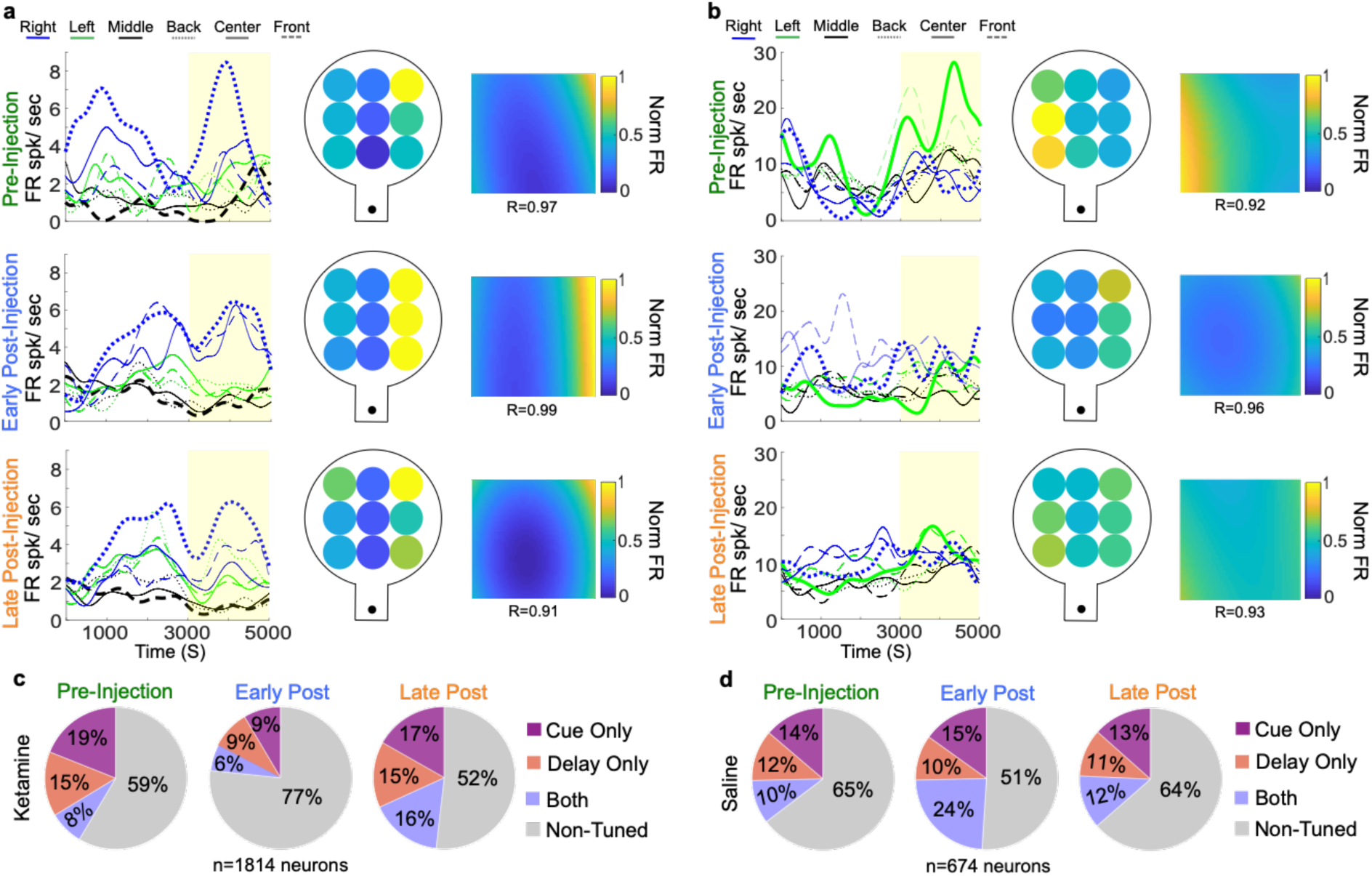
Tuning of single neurons for memorized locations and effects of ketamine. **a**, Firing rate of an example neuron for a ketamine-WM session. On the left, SDFs over cue and delay (yellow) epochs. Preferred locations and least-preferred locations are bolded. Center, firing rates during the delay epoch for all target locations. Right, firing rates fitted to a polynomial plane. **b**, Firing rate of a second example neuron during a ketamine-WM session. **c**, Average proportion of tuned single units during the cue epoch (pink), delay epoch (orange), or during both (purple) for each injection period for ketamine-WM sessions. **d**, Average proportion of tuned single units during each epoch for saline-WM sessions.

### Population decoding

Single neuron tuning is essential for information coding. However, the information encoded by a neuronal population can only be evaluated by examining the activity of simultaneously recorded neurons^4,23^. We used a linear classifier (Support Vector Machine, SVM) to predict from neuronal ensemble activity whether targets were presented on the left, right or center of the virtual arena on a single trial basis. We pooled locations in order to reach a sufficient sample size (trials) to use cross-validation procedures. Decoding accuracy for different ensemble sizes was higher than chance (33%) in all analyzed experimental sessions (Fig. 3a, b and Extended Data Fig. 3). Decoding accuracy decreased after ketamine injection between pre-injection and post-injection periods (Fig. 3a), predominantly during the delay and response epochs. The classifier made systematically more errors after ketamine injection, as animals did. Similar results were observed when using only correct trials or decoding 9 target locations in sessions with sufficient sample sizes (Extended Data Fig. 4). On the other hand, decoding accuracy remained stable between injection periods in saline-WM sessions (Fig. 3b). These results indicate that LPFC neuronal ensembles encode spatial WM in naturalistic settings and that ketamine disrupts these ensemble codes.

**Figure 3.**
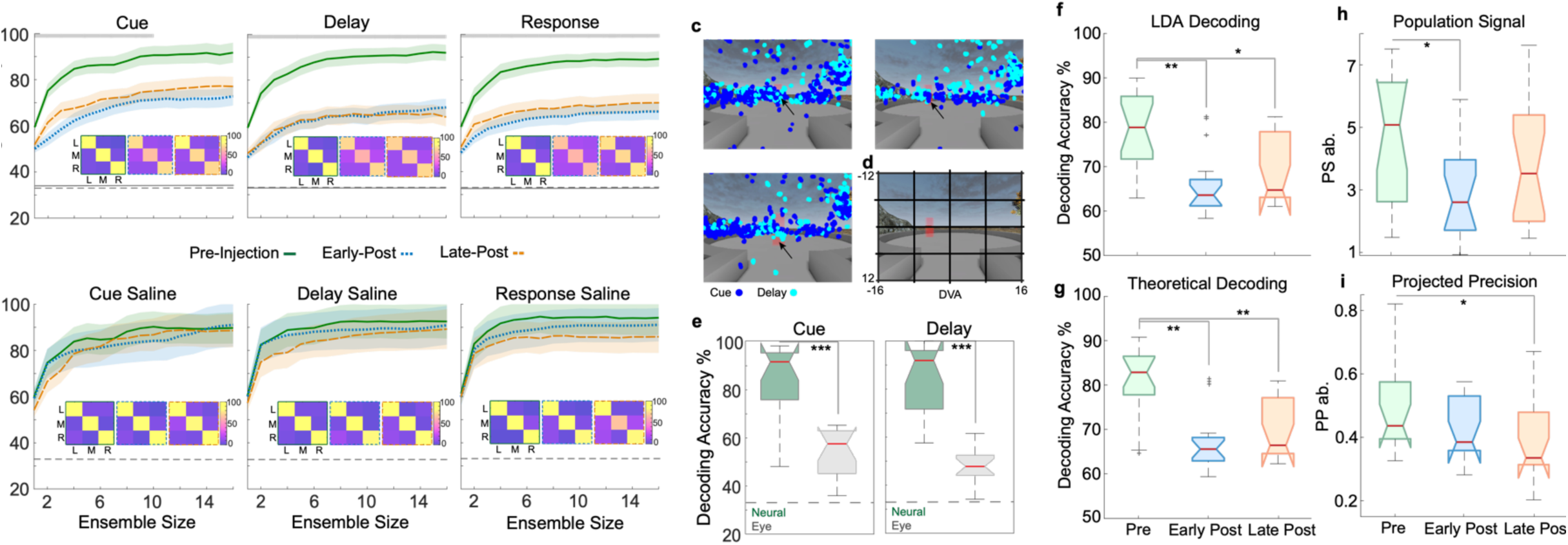
Population analyses. **a**, Median decoding accuracy for ketamine-WM sessions for pre-injection (green), early post-injection (blue), and lat post-injection periods (orange) for trial epochs. Chance performance is indicated by dashed grey line and shuffled result are indicated by solid grey line. Confusion matrices for each injection period indicate classifier performance for each target location. Grey bars near top of the plot indicate ensemble sizes showing a significant reduction in decoding accuracy from pre-injection to early post-injection periods (Kruskal Wallis, p<0.05). **b**, Same as **a**, for saline-WM sessions. Al error bars are SEM. **c**, Fixation locations on screen for an example session during the cue and delay epochs for three targe locations indicated by arrows. **d**, Screen divided into 16 regional cells. **e**, Comparison between decoding target locatio accuracy using neuronal ensemble activity (green) and eye fixation position on screen (grey) during the pre-ketamin injection period for the cue and delay epochs. **f**, DPe over the delay epoch for ketamine-WM sessions. **g**, DPt over th delay epoch for ketamine-WM sessions. **h**, PS over the delay epoch for ketamine-WM sessions. **i**, PP over the delay epoch for ketamine-WM sessions. Red center lines indicate median, the bottom and top edges of the box indicate the 25th and 75th percentiles. The whiskers extend to non-outlier data points (approximately within 2.7 std) and the outliers are plotted using ‘+’. *<0.05, **<0.01, ***<0.001.

### Gaze decoding

A proportion of neurons in the LPFC encode signals related to gaze^11^. Since gaze was unconstrained in our task, it is possible that the coding of remembered locations predominantly reflect systematic biases in eye position signals. To explore this possibility, we first determined whether animals showed biases in eye position towards the target location. We used a linear classifier to predict target location from the position of eye fixations on the screen. We divided the screen into 16 cells (see Fig. 3d) and calculated the number of fixations falling within each cell. During the pre-injection period, the accuracy for decoding remembered locations from fixations was significantly higher than chance, indicating a target specific gaze bias (cue: T-Test, *p*<0.0001, delay: *p*<0.0001; Fig. 3e). Such a bias was less pronounced during the delay relative to the cue epoch (Wilcoxon Rank-sum, *p*=0.002; Extended Data Fig. 5a, b). However, decoding accuracy for remembered locations from eye position was significantly lower than decoding accuracy of a classifier that uses neuronal firing rate and the same number of features (n=16) (Kruskal Wallis, cue: *p*=0.0001; delay: *p*<0.0001; Fig. 3e). This suggests that biases in eye position signals are not sufficient to account for the amount of information encoded by the population activity regarding target location.

Decoding accuracy for eye position remained stable after ketamine injection indicating that low doses of ketamine did not significantly affect the documented biases in gaze position (Kruskal Wallis; cue, *p*=0.135, delay, *p*=0.101; Extended Data Fig. 5a, b). On the other hand, decoding accuracy from neuronal activity significantly decreased after ketamine injection (Fig. 3a). These results indicate that biases in eye position cannot account for the effects of ketamine on decoding of target locations from neuronal activity and suggest a dissociation between eye position and WM signals within LPFC microcircuits.

Finally, we decoded eye position on screen from neuronal firing rates during eye fixation periods. Neither the decoding accuracy for the cue or delay epoch significantly differed from chance (T-Test, cue: *p*=0.117, delay: *p*=0.646) (Extended Data Fig. 5d), indicating that in our naturalistic task, representations of the target location were dissociated from eye position signals. Together, these results illustrate that population codes for spatial WM in the LPFC are dissociable from changes in retinal signals and saccadic eye movements.

### Statistical properties of neural decoding

The amount of information encoded by a population of neurons is determined by two core statistical properties of population activity, the population signal (PS) and the projected precision (PP)^23^. PS reflects differences in neurons’ individual tuning and the modulation of the population response across target locations (i.e. the length of the vector between population responses to different target locations). PP reflects the trial to trial variability of neuron activity and accounts for the projection of the covariance matrix inverse on the direction of the PS vector. It is possible that the effect of ketamine on neural coding occurs through modulation of one or both properties. To investigate this, we analyzed the effects of ketamine on the ability of neuronal populations to discriminate remembered locations on the left or right side of the arena. We used binary classes (left vs. right locations) and ensembles of 3 neurons in order to reach a large enough sample size to reliably compute PS and PP^23^. We selected random ensembles providing the highest empirical decoding accuracies for the remembered location (within the top 75th percentile calculated using Linear Discriminant Analysis, LDA).

We first compared decoding accuracy results using empirical decoding (LDA) and theoretical decoding (using PP and PS) and demonstrate no significant differences (Kruskal Wallis, *p*=0.171) (Extended Data Fig. 6a, b). This shows that our theoretical decoding method employing PP and PS accurately estimates information content of neural ensembles.

As shown in Fig. 3f and Fig. 3g, there was a significant decrease in empirical decoding accuracy (DPe; Kruskal-Wallis, post hoc, *p*=0.001) and theoretical decoding accuracy (DPt) (Kruskal-Wallis, post hoc, *p*=0.0002) after ketamine injection compared to the pre-injection period and an increase during the late post-injection compared to the early post-injection period (for data per NHP see Extended Data Fig. 6c-f). Decoding accuracy in saline-WM sessions did not significantly change between injection periods (see Extended Data Fig. 6g, h). Notably, PS significantly decreased after ketamine injection compared to the pre-injection period (Kruskal-Wallis, post hoc, *p*=0.012) and increased during the late post-injection compared to the early post-injection period (Fig. 3h). Contrarily, PP showed a small non-significant decrease in the early post-injection period compared to the pre-injection period (Kruskal-Wallis, post hoc, *p*=0.38) (Fig. 3i) (for data per NHP see Extended Data Fig. 6i-l). The PP drop became significant during late post-injection relative to pre-injection (p=0.012). The latter result may suggest that ketamine induced slow changes in correlated variability or its projection onto the PS vector which outlasted changes in neuronal tuning. Overall these results indicate that the observed early changes in information decoded from populations of neurons after ketamine injection is primarily due to changes in PS, a consequence of changes in individual neuron tuning.

### Cell type specific effects of ketamine

Ketamine induces a variety of effects on individual neurons^14,24^. A loss of neuronal tuning may result from neurons increasing their response to least-preferred locations (see example neuron Fig. 2a) or decreasing their response to preferred locations (see example neuron Fig. 2b). One possible explanation for this heterogeneity is that different cell types (e.g., pyramidal cells and interneurons) may be differentially affected by ketamine. To test this hypothesis, we divided neurons into narrow and broad spiking based on waveform peak-to-trough duration or width (Fig. 4a, b). In mouse neocortex, broad spiking neurons are largely putative pyramidal cells or in smaller proportion, vasointestinal peptide expressing (VIP) neurons. On the other hand, narrow spiking neurons are largely parvalbumin (PV) expressing, or in a smaller proportion, somatostatin (SST) expressing neurons^25^.

**Figure 4.**
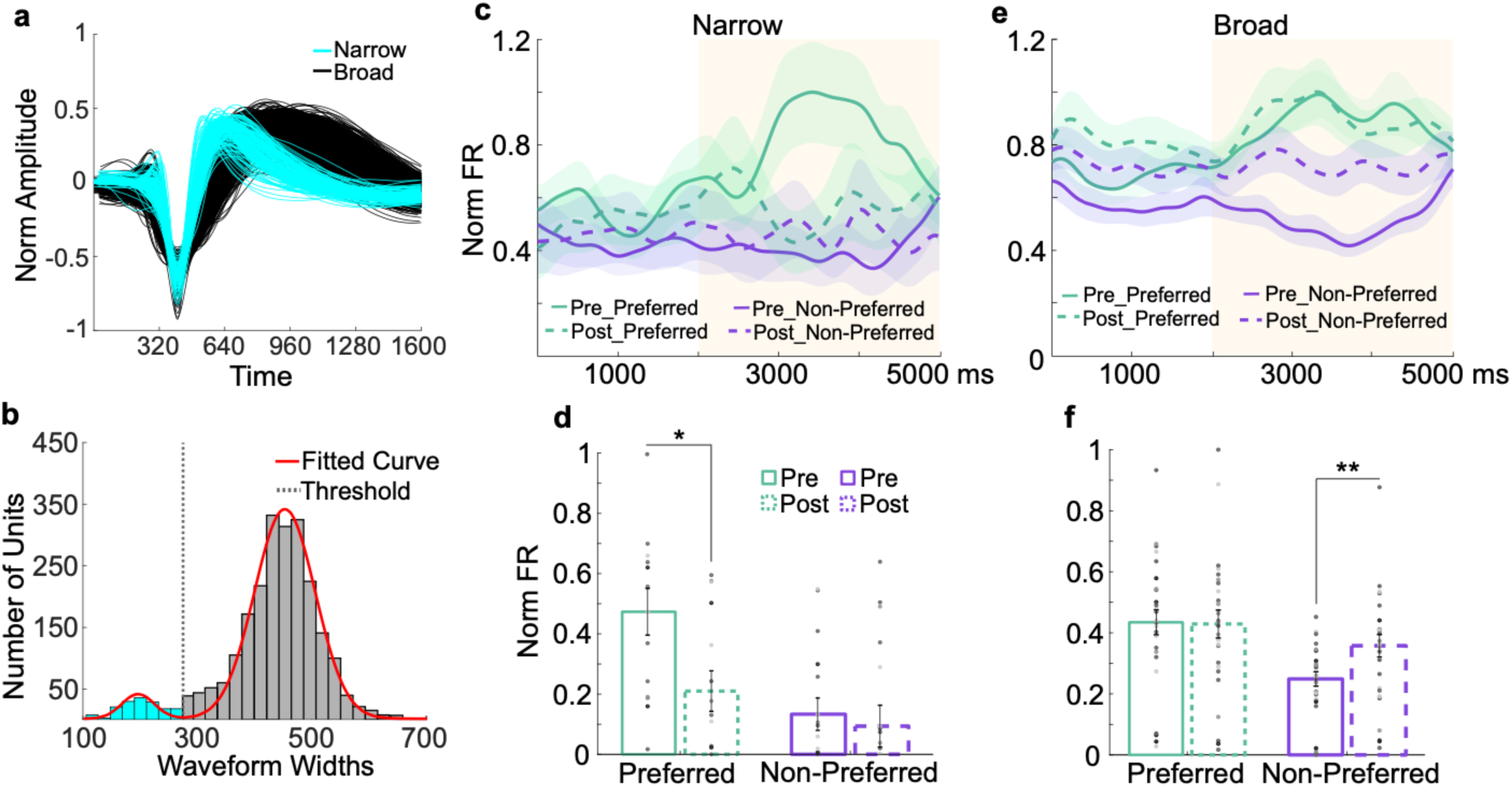
Cell type specific effects of ketamine on working memory signals. **a**, Waveforms of narrow and broad spiking neurons. **b**, Distribution of waveform widths (microseconds) fitted with a 2-Gaussian model. Boundary line between narrow and broad spiking neurons is at the intersection point between Gaussians (275, dotted line). Gaussian at the lower width boundary indicates narrow spiking neurons (blue) and the upper boundary indicates broad spiking neurons (dark grey). **c**, Normalized average population SDFs for cue and delay (yellow) epochs for delay tuned narrow spiking neurons. **d**, Median population SDF for narrow spiking neurons over the delay epoch. Data points represent value per electrode array for each session. **e**, Normalized average population SDF for cue and delay epochs (yellow) for delay tuned broad spiking neurons. **f**, Median population SDF for broad spiking neurons over the delay epoch. All error bars are SEM. *<0.05, **<0.01, ***<0.001.

After ketamine injection, narrow spiking neurons show a loss of tuning during the delay epoch due to a decrease in firing for their preferred locations compared to the pre-injection period (Wilcoxon Rank-sum, *p*=0.049) with no significant change for their least-preferred locations (*p*=0.546) (Fig. 4c, d). In contrast, broad spiking neurons show a loss of tuning due to a significant increase in firing for their least-preferred locations compared to the pre-injection period (Wilcoxon Rank-sum, *p*=0.006) with no significant change for their preferred locations (*p*=0.649) (Fig. 4e, f). Such changes were not observed during saline-WM sessions (Extended Data Fig. 7a, b; data per NHP in Extended Data Fig. 7c-j). We also conducted separate analyses of PS in narrow and broad spiking single neurons and found a loss of PS and a resultant decrease in DPt in both populations (see Extended Data Fig. 8).

Considering that our populations of NS and BS neurons are dominated by PV and pyramidal cells respectively, our findings align with a proposed pathophysiological mechanism for WM dysfunction: reduced NMDAR conductance on PV interneurons, amounting to generalized disinhibition of pyramidal cells and resultant loss of tuning^24, 26^. Indeed, ketamine has high affinity for GluN2B NMDAR subunits which are expressed in PV interneurons^27,28^. Loss of pyramidal cells tuning reduces the spatial specificity of WM representations, the PS, and encoded information regarding remembered target location.

## Conclusion

Our study shows that macaque LPFC neurons encode WM representations during naturalistic tasks, regardless of potential interference by sensory and motor signals generated during natural behavior. The LPFC differs from other areas such as the posterior parietal cortex where WM representations are perturbed by visual distractors^7^. The emergence of the granular LPFC allows for the encoding of representations that are dissociated from distraction and action. Such an emergence occurred in anthropoid primates, expanding their mental world and consequently enhancing their adaptability to changing environments^6,29^. Moreover, the observed effects of ketamine indicate that mental codes in LPFC rely on a delicate balance between the activation of excitatory and inhibitory neuronal types mediated by NMDA receptors. A break-down of this balance may explain cognitive symptoms found in schizophrenia and other brain diseases exhibiting LPFC abnormalities and NMDAR hypoactivity^13,17,30^.

## Methods

Two adult male rhesus macaques (*Macaca mulatta*) were used in this experiment (age: 10, 9; weight: 12, 10 kg). Results shown in the main text and figures represent results across subjects unless otherwise specified.

### Ethics statement

Animal care and handling including basic care, animal training, surgical procedures, and experimental injections were pre-approved by the University of Western Ontario Animal Care Committee. This approval ensures that federal (Canadian Council on Animal Care), provincial (Ontario Animals in Research Act), regulatory bodies (e.g: CIHR/NSERC), and other national CALAM standards for the ethical use of animals are followed. Regular assessments for physical and psychological wellbeing of the animals were conducted by researchers, registered veterinary technicians, and veterinarians.

### Task

The current task takes place in a virtual environment. This environment was developed using Unreal Engine 3 development kit: utilizing Kismet sequencing and UnrealScript (UDK, May 2012 release; Epic Games). More about this platform and the recording setup can be found in Doucet, Gulli, and Martinez-Trujillo, 2016. Within this virtual environment, target locations were arranged in a 3 × 3 grid, spaced 290 unreal units apart (time between adjacent targets is approximately 0.5 seconds). Movement speed was fixed throughout navigation.

### Experimental setup

The task was presented on a computer LDC monitor positioned 80 cm from the subjects’ eyes (27” ASUS, VG278H monitor, 1024 × 768 pixel resolution, 75 Hz refresh rate, screen height equals 33.5 cm, screen width equals 45 cm). Subjects performed the experiment in an isolated room with no illumination other than the monitor. The walls, doors, and ceiling of the room were RF shielded and contained no AC power lines. Cables providing power to the setup equipment entered the room through a small aperture in a wall and were shielded to minimize interference with the recordings. Eye positions were monitored using a video-oculography system with sampling at 500 Hz (EyeLink 1000, SR Research). A custom computer program-controlled the stimulus presentation (through Unreal Engine 3), reward dispensation, and recorded eye position signals and behavioral responses. Subjects performed the experiment while seated in a standard enclosed primate chair (Neuronitek) and were delivered juice reward through a tube attached to the chair and an electronic reward integration system (Crist Instruments). Prior to the experiments, subjects were implanted with custom fit, PEEK cranial implants which housed the head posts and recording equipment (Neuronitek). See Blonde et al, 2018 for more information. The head posts attached to a head holder to fix the monkey’s heads to the primate chair during training and experimental sessions.

### Microelectrode array implant

Surgical procedures were conducted under general anesthesia induced by ketamine and maintained using isoflurane and propofol. Two 10×10, microelectrode Utah arrays (96 channel, 1.5 mm in length, separated by at least 0.4 mm) (Blackrock Microsystems) were chronically implanted in each animal. They were located in the left LPFC — anterior to the arcuate sulcus and on either side of the posterior end of the principal sulcus^12^. Brain navigation for surgical planning was conducted using Brainsight (Rogue Research Inc.) (see Extended Data Fig 9a, b). Arrays were placed and impacted approximately 1.5 mm into the cortex. Reference wires were placed beneath the dura and a grounding wire was attached between screws in contact with the pedestal and the border of the craniotomy. Electrode placement was approximated using CT imagining post-operatively (Extended Data Fig. 9c).

### Neural recordings and spike detection

Neural data was recorded using a Cerebus Neuronal Signal Processor (Blackrock Microsystems) via a Cereport adapter. The neuronal signal was digitized (16 bit) at a sample rate of 30 kHz. Spike waveforms were detected online by thresholding at 3.4 standard deviations of the signal. The extracted spikes were semi-automatically resorted with techniques utilizing Plexon Offline Sorter (Plexon Inc.). Sorting results were then manually supervised. Multiunits consisted of threshold-crossing events from multiple neurons, with action potential-like morphology, that were not isolated well enough to be classified as a well-defined single unit (for spike sorting example see Extended Data Fig. 9d, e). We collected behavioural data across 18 ketamine-WM sessions (9 in NHP T, 9 in NHP B) and neural data from 17 ketamine-WM sessions with one session from NHP T removed due to incomplete synchronization of neural data during the recording. This yielded a total of 2906 units recorded during ketamine-WM sessions: 1814 single neurons (259 in NHP T, 1555 in NHP B) and 1092 multiunits (533 in NHP T, 559 in NHP B). Behaviour and neural data was recorded from seven saline-WM sessions resulting in 1117 units in total: 674 single units (48 in NHP T, 626 in NHP B) 443 multiunits (126 in NHP T, 317 in NHP B). Behavioural data from four ketamine-perception sessions were analyzed (2 in NHP T, 2 in NHP B).

### Ketamine injection

Animals were trained to voluntarily receive injections in the primate chair while in the experimental setup. An intramuscular injection of either ketamine (0.25, 0.4, or 0.8 mg/kg) or saline (0.25 mg/kg) was administered in the hamstring muscles by a registered veterinary technician. The ketamine doses were titrated so they did not induce visible behavioral changes in the animals such as nystagmus or somnolence. Ketamine injections were spaced at least two days apart to allow for washout of the drug^33^.

### Behavioural analysis

Correct trials are trials in which subjects reach the correct target location within 10 seconds. The percent of correct trials was compared to chance (11%) for each session using binomial tests.

The percent of correct trials over time was calculated using 15 equally sized trial bins for each injection period. The resulting 45 data points per session were averaged over all ketamine-WM and saline-WM sessions for each animal and then combined across subjects. Statistical analysis was conducted by comparing the percent of correct trials binned over the three injection periods (pre, early post, and late post-injection periods) for ketamine-WM and saline-WM sessions. Response time was calculated for correct trials as the duration between navigation onset and end of trial for each experimental condition (target location) for each recording session.

### Trajectory analysis

Analyses of an animal’s trajectories within the navigation period are conducted on trials in which the animal crosses a predetermined line that divides the start enclave from the main body of the task arena. The task environment was divided into a 5×5 grid containing 25 regional cells of equal dimensions. The grid was sized so that cells tightly enclose target locations (see Fig. 1j). For each trial, we calculated whether the subject entered each cell during navigation resulting in either 1 (entered) or 0 (not entered) per cell. The number of trials in which each cell was entered was then divided by the total number of trials. This resulted in a percent value for each cell for each target location condition (25*9 conditions, n=225 per session) that was then averaged over all ketamine-WM or saline-WM sessions. We then calculated increases in average percent values for each cell between injection periods (values above 0 included).

### Spatial selectivity

Single units (2488 from 17 ketamine-WM and seven saline-WM sessions) were tested for selectivity for target location during a given epoch for correct trials by computing a one-way analysis of variance on epoch-averaged firing rates with target location as the independent variable. A unit was defined as selective if the test resulted in *p* < 0.05. A neuron’s preferred location was defined as the location that elicited the largest response during the epoch of interest. The least-preferred location was defined as the location that elicited the smallest response. The proportion of tuned single units for each task epoch (cue and delay) were compared between injection periods for ketamine-WM and saline-WM sessions.

### Spike density functions

The activity of tuned single neurons were plotted as trial averaged spike density functions for each task condition (target location) which were generated by convolving the spike train with a Gaussian kernel (standard deviation=150 ms). We normalized responses by the maximum firing rate in each neuron’s preferred target location.

### Plane Fitting

In order to visualize neural responses to different target locations within the 2D space, we fit a second order polynomial surface to the mean normalized firing rate for the 9 target location conditions to the x- and y-coordinates of each target location. Firing rate was normalized by the maximum firing rate in the pre-ketamine injection period. This method was used for visualization (Fig. 2a, b), not for quantitative analysis.

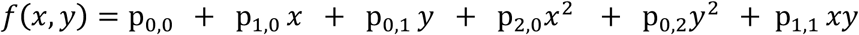

### Neuronal ensemble decoding

We used a linear support vector machine (SVM) (Libsvm 3.14)^34^ with 5-fold cross-validation to extract task-related activity from z-score normalized population-level responses using both single units and multiunits on a single trial basis. The regularization parameter used was the optimal penalty parameter *C* (refer to Eq. 1 in Fan et al. 2008). The classifiers used firing rates calculated over epoch durations (cue, 3000ms; delay, 2000ms; response first 2000ms) from ensembles of neurons simultaneously recorded within each session to predict target location for correct and incorrect trials within the virtual arena (left, center, right).

For each behavioral session, we calculated decoding performance for neuronal ensembles with a maximum of 16 neurons since decoding performance plateaued around this point (see Extended Data Fig. 3a, b). We began building ensembles by selecting the neuron with highest individual performance in decoding target location. This neuron was then paired with all remaining neurons to find the pair of neurons that maximized decoding performance. We then used this pair and combined it iteratively with all remaining neurons to find the best trio. This procedure was repeated until 16 neurons were reached.

We pooled target locations across depth in order to have a sufficient number of trials for training and testing the classifiers. We chose to combine trials based on target direction in the environment (left, center, right) based on observations that neurons tended to show more similar responses to targets located in the same direction compared to targets located at the same depth within the environment. Observations were balanced between classes using subsampling (without replacement) which was repeated 20 times.

We maintained the same neurons in ensembles (for ensembles of 16 neurons) and used the same procedure to calculate chance performance obtained by randomizing class labels (all other data features remained unaltered). We repeated this shuffling procedure 10 times for each session. Subsampling was conducted 20 times in each iteration. Using this procedure, the shuffled decoding accuracy for one ketamine-WM session from NHP T was higher than expected by chance; therefore, this session was removed from the analysis. Decoding accuracy between injection periods for ketamine-WM and saline-WM sessions was compared for each neuronal ensemble size. We ran the decoding procedure a second time restricting to correct trials only (in sessions with a sufficient number of samples for cross-validation). Finally, a third decoding analysis was conducted using all 9 target locations from neural data on a single trial basis using SVM with 4-fold cross validation (in sessions with a sufficient number of samples: 1 session in NHP T, 2 sessions in NHP B).

### Empirical decoding

Classification was performed using Linear Discriminant Analysis (LDA) with regularization on 1000 random neural ensembles of two, three, and five units using single units and multiunits (with firing rates >0.5 Hz). Decoding accuracy was determined using 5-fold cross-validation. Since this analysis depends on binary data, trials (based on target locations) were grouped as being presented on the right or left of the environment (chance=50%). Trials in which centrally placed targets are presented are not used in this analysis.

Further analysis was conducted using the top performing ensembles for each injection period (ensembles with the highest information content, decoding accuracy in the 75th percentile).

### Theoretical decoding accuracy

Theoretical decoding accuracy (DPt) was calculated using the same 1000 random neural ensembles of two, three, and five units as the empirical decoding analysis using previously described statistical properties of the population response^23^. DPt was calculated as:

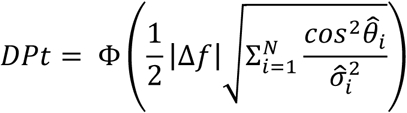

where Φ represents cumulative Gaussian of trial firing rates, the first term, |Δ*f* | represents the Population Signal (PS) that measures target condition specific modulation of the population response (population tuning) and the second term,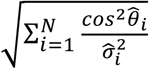, represents projected precision (PP), which is a function of 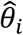, the angle between the i-th eigenvector of the covariance matrix Σ and the direction of the stimulus 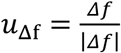 vector tuning, and 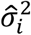 the i-th eigenvalue of the covariance matrix (see supplemental material for illustration). PP measures the population response variability (trial to trial variability). Further analysis was conducted using top performing ensembles (as classified by empirical decoding using LDA).

### Waveform classification

Single units were classified as either narrow (NS) or broad spiking (BS) based on action potential width measured as peak-to-trough interval duration^25^. Average waveforms for each unit were interpolated with a cubic spline fit to increase the resolution of the data. The duration between waveform peak and trough were then calculated based on time stamps from the minimal and maximal voltage values. Waveform widths for all neurons were plotted in a histogram. After removing outlier widths (>675 microseconds), 2314 units remained and are included in the analysis. A bimodal distribution was visualized and then quantified by fitting the data with either a single (1-Gaussian) or sum of two Gaussian functions (2-Gaussian) to determine optimal fit. The goodness of fit for both functions was determined using Akaike Information Criterion (AIC)^35^ with the lowest value determined for 2-Gaussians indicating bimodality.

The threshold dividing NS and BS (275 microseconds) was determined by setting a boundary at the inflection point of the two Gaussian fitted distributions (Fig. 4b)^25, 36^. Waveform amplitudes were normalized to the difference between highest and lowest amplitudes for each unit waveform and waveforms were aligned at threshold crossing for visualization (Fig. 4a). Based on this threshold, 161 neurons were classified as NS and 2153 neurons were classified as BS. 750 tuned broad spiking neurons were included for further analysis for ketamine-WM sessions and 246 tuned units were included for saline-WM sessions. 41 tuned narrow spiking neurons were included for ketamine-WM sessions and 11 tuned neurons were included for saline-WM sessions.

### Firing rate for preferred and non-preferred locations

Spike density functions (SDF) were calculated for NS and BS neurons that were significantly tuned for remembered locations during the delay period (ANOVA, *p*<0.1). We specifically obtained the spike density functions for these neurons for their preferred and least-preferred locations during the cue and delay epochs before and after ketamine or saline injection. Population activity was calculated by averaging SDFs between simultaneously recorded single units within the same array and responses were normalized by the maximum population response. These population responses for each array were then averaged over all ketamine-WM or saline-WM sessions. Firing rates were averaged during the delay epoch and were statistically compared using 1-tailed Wilcoxon Rank-sum tests between pre and early post-injection periods for preferred and least-preferred locations.

### Gaze analysis

Gaze position was computed from eye tracking signals synchronized with the neural recordings and behavioral performance measurements^37^. The amount of time that gaze fell within the screen boundaries was calculated during the cue and delay epochs of the task and were statistically compared before and after ketamine or saline injection (Extended Data Fig. 10a-f).

Eye movements were classified as saccades, fixations, or smooth pursuits based on previously published methods for eye movement classification in virtual environments in which periods of high acceleration approximate saccade epochs and movement patterns were used to determine precise saccade onset and offset. Foveations are classified as fixations or smooth pursuits based on measures of spatial range (see Corrigan, Gulli, Doucet, & Martinez-Trujillo, 2017 for detailed method). The proportion of fixations falling within the trial specific target location compared to other potential target locations on the screen was calculated (Extended Data Fig. 10g-l).

### Decoding eye position

During the cue and delay epochs, the screen was divided into 16 cells of equal dimensions. The number of foveations classified as fixations were calculated within each cell under the assumption animals gather information from the virtual environment during such fixation periods^37^. We used a linear classifier (SVM) with 5-fold cross-validation to determine whether target location could be predicted on a single trial basis by the number of fixations within each cell: the extent to which animals fixate in each part of the visual environment. This analysis was compared with a decoding analysis using neural ensembles utilizing the same number of features (16 neuron ensembles).

### Decoding eye position and target condition from neural data

We used a linear classifier (SVM) with 4-fold cross validation to decode eye position on screen based on neural firing rates during fixations. Four target locations were selected as part of this analysis since their location onscreen were easily separable. Four regions on the screen were outlined surrounding these target locations. Fixation periods occurring in either the cue or delay epoch that fell within these regions were used (see Extended Data Fig. 5c). Short fixation periods were removed (amplitude < 6 ms). Firing rate was calculated for each neuron during each fixation period and were z-score normalized. Sessions missing observations (fixation periods) for 1 or more classes were excluded from this analysis.

## Acknowledgements

We thank registered veterinary technicians Kim Thomaes and Rhonda Kersten from the University of Western Ontario for their assistance in surgery and animal care; Guillaume Doucet from the University of Ottawa for technical assistance related to Unreal Development Kit; Maryam Nouri Kadijani from the University of Western Ontario for assisting with initial data exploration; Kevin Barker from Neuronitek for engineering equipment for our experiments; Jonathan C. Lau from the Division of Neurosurgery, University Hospital for providing advice regarding surgery and surgical planning; Matthew Leavitt, AI Resident at Facebook for access to MATLAB code related to polynomial plane fitting and advice on electrophysiological analysis. This work was supported by Canadian Institute of Health Research Project Grant; Natural Sciences and Engineering Research Council of Canada (NSERC); Ontario Graduate Scholarship; Jonathan & Joshua Memorial Graduate Scholarship in Mental Health Research. Chrysalis Foundation (London, Ontario). L.P. acknowledges salary support from the Tanna Schulich Endowment Chair for Neuroscience and Mental Health. R.M.-B. acknowledges support from MINECO (Spain; BFU2017-85936-P), the Howard Hughes Medical Institute (HHMI, ref 55008742), and the ICREA Academia (2016),

## Author Contributions

M.R., L.P., and J.C.M.-T. planned the study. M.R. and R.L. trained animals, performed the experiments and conducted data preprocessing including spike sorting. M.R. analyzed the data, created figures and wrote the manuscript with help from J.C.M.-T. L.R.D. developed unique MATLAB code for several analyses and contributed knowledge regarding machine learning. B.C. developed unique MATLAB code for eye movement classification and analysis. R.A.G. developed code for data preprocessing and contributed essential knowledge of experimental design and data analysis. B.C. and R.A.G. trained animals to perform eye fixations for eye tracking calibration. R.N. and R.M.-B. contributed to analysis design. A.J.S., J.C.M.-T., R.A.G., R.L., & and M.R. planned and conducted surgeries.

## Data availability

Data supporting the findings of this study is available from the corresponding authors on reasonable request and will be fulfilled by M.R.

## Code availability

MATLAB codes used the analyze the data are available from M.R.

**Supplementary Materials: Online Movie 1. Working Memory Task**.

https://www.youtube.com/watch?v=nZDYJw2aFLQ

One example trial of the working memory task. Featured: Cue, delay, and response epochs. Monkey’s eye position in the virtual environment are indicated by the moving white circle with label of objects that falls within foveated position.

**Supplementary Materials: Online Movie 2. Ketamine’s Effect on Task Performance**.

https://youtu.be/r5ouvtSx_XQ

Example trials of the working memory task before and after ketamine injection. Featured: Cue, delay, and response epochs. Monkey’s eye position in the virtual environment are indicated by the moving white circle with label of objects that falls within foveated position.

## Extended Data

**Figure 1.**
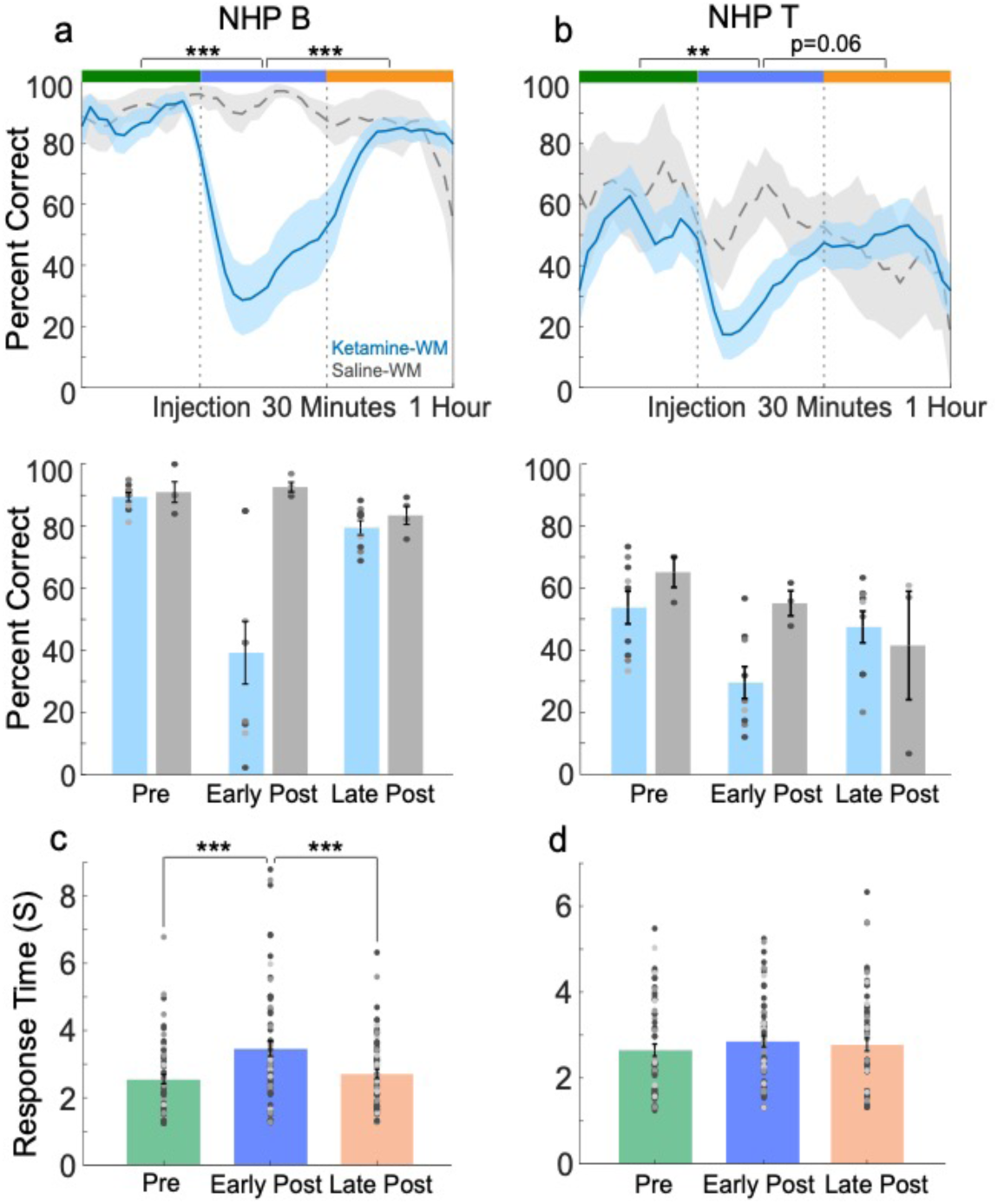
Task performance per subject. **a**, Average percent of correct trials over ketamine-WM (blue) and saline-WM (grey) sessions for NHP B. **b**, Average percent of correct trials for ketamine-WM (blue) and saline-WM (grey) sessions for NHP T. **c**, Average response time for correct trials for NHP B for each injection period. **d**, Average response time for correct trials for NHP T for each injection period. All error bars are SEM. *<0.05, **<0.01, ***<0.001.

**Figure 2.**
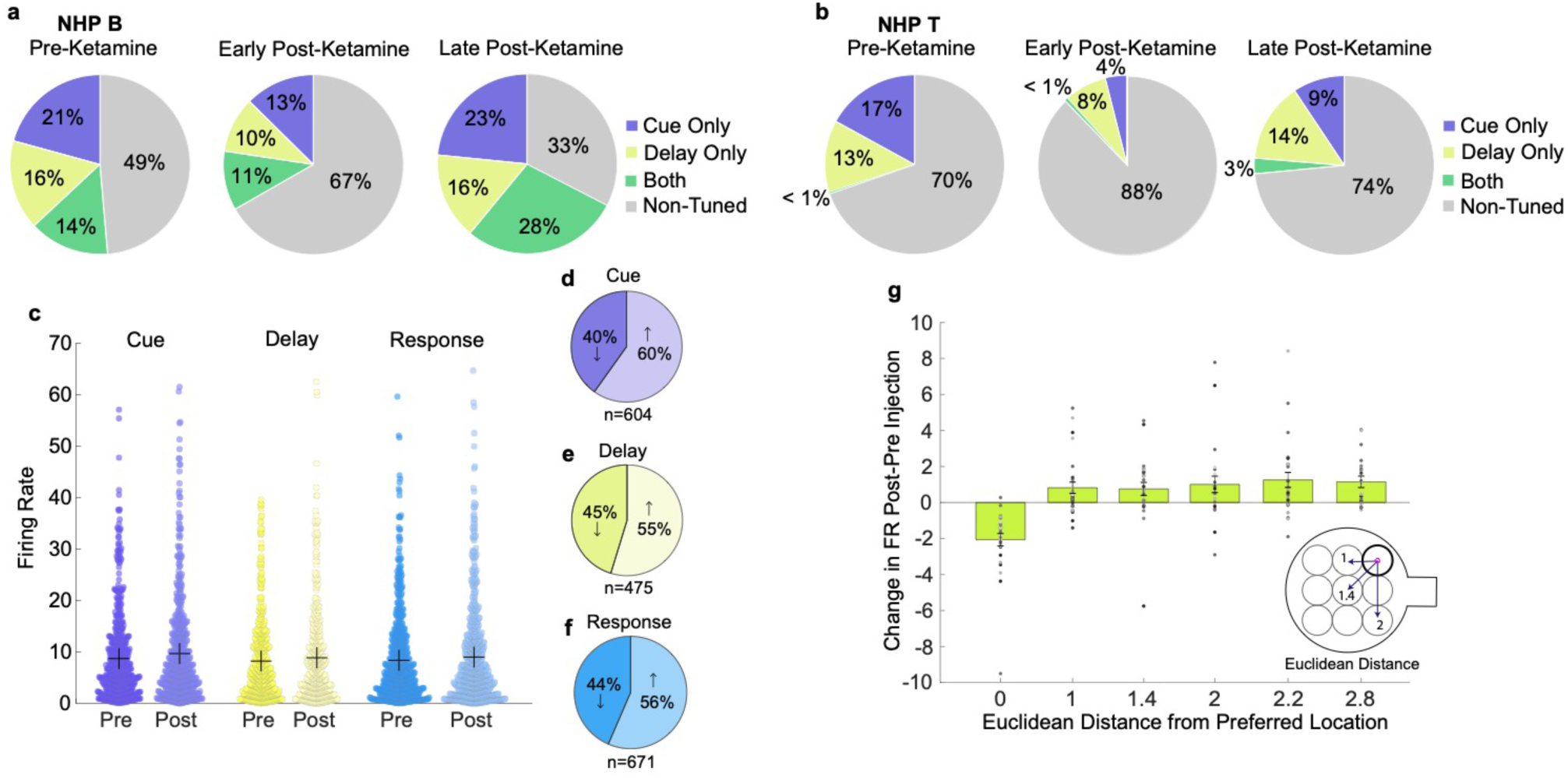
Changes in neuron tuning and firing rate. **a**, Average proportion of tuned single units for ketamine-WM sessions for NHP B during the cue, delay, or both epochs. **b**, Average proportion of tuned single units for ketamine-WM sessions for NHP T for the trial epochs. **c**, Difference in firing rate of single units for correct trials during cue, delay, and response epochs between pre and early post-injection periods for ketamine-WM sessions. Mean indicated by ‘+’. **d**, Proportion of cue tuned single units showing either an increase or decrease in firing rate after ketamine injection during the cue epoch for correct trials. **e**, Proportion of delay tuned single units showing either an increase or decrease in firing rate after ketamine injection during the delay epoch for correct trials. **f**, Proportion of response tuned single units showing either an increase or decrease in firing rate after ketamine injection during the first 2000 ms of the response epoch for correct trials. **g**, Average change in firing rate for single units from the pre-injection to the early post-injection period as a function of Euclidean distance from neuron’s original preferred location during the delay epoch for each target location for ketamine-WM sessions. Euclidean distance of 0 indicates preferred location and data points represent the average of each array for each session. All error bars are SEM. *<0.05, **<0.01, ***<0.001.

**Figure 3.**
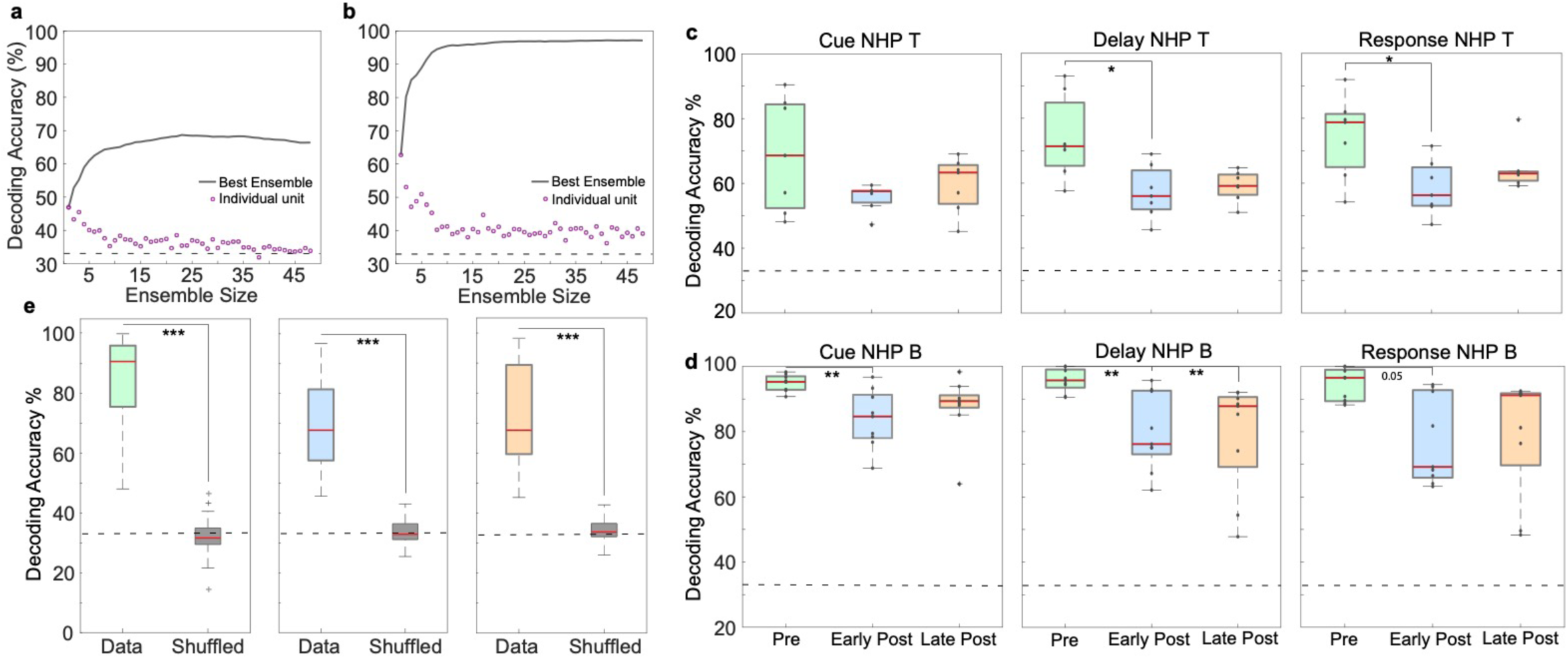
Decoding performance per subject. **a**, Single NHP T session example of decoding accuracy as a function of neuronal ensemble size. **b**, Single NHP B session example of decoding accuracy as a function of neuronal ensemble size. Purple data points represent individual neuron contribution to decoding accuracy. **c**, Sixteen neuron ensemble decoding accuracy for trial epochs over injection periods for ketamine-WM sessions for NHP T. **d**, Sixteen neuron ensemble decoding accuracy for trial epochs over injection periods for ketamine-WM sessions for NHP B. Data points represent decoding accuracy per session. **e**, Decoding accuracy for ketamine-WM sessions with data combined between trial epochs for injection periods. Decoding accuracy for shuffled data in grey (shuffled trial condition labels). Red center lines indicate median, the bottom and top edges of the box indicate the 25th and 75th percentiles. The whiskers extend to non-outlier data points (approximately within 2.7 std) and the outliers are plotted using ‘+’. *<0.05, **<0.01, ***<0.001.

**Figure 4.**
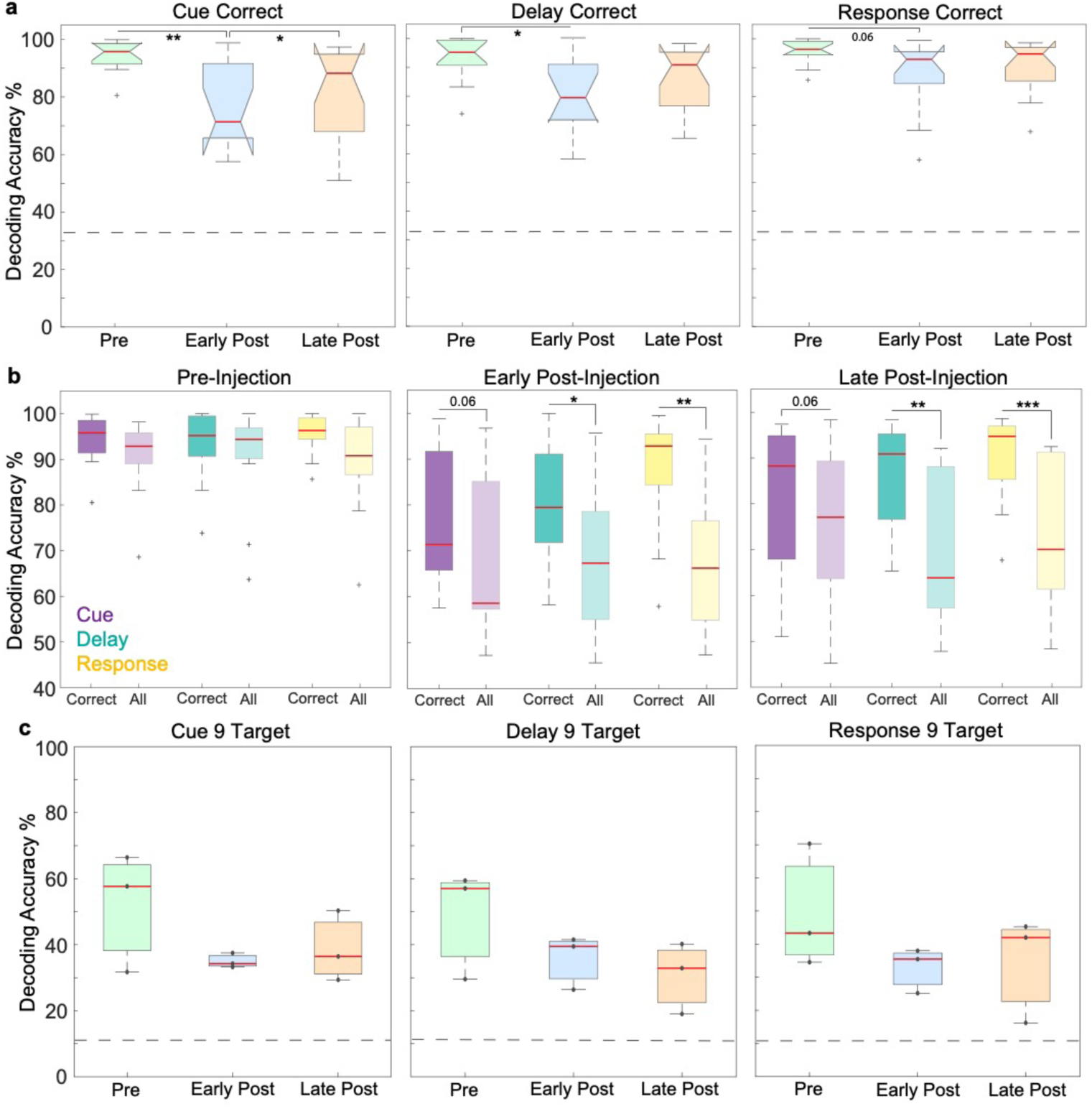
Ensemble decoding for correct trials and 9 target locations. **a**, Decoding target location from neuronal ensembles using correct trials. Decoding accuracy for ketamine-WM sessions for pre-injection (green), early post-injection (blue), and late post-injection periods (orange) for trial epochs. Chance performance is indicated by dashed grey line. **b**, Comparison between decoding accuracy using correct trials and using all trials for trial epochs and injection periods. **c**, Decoding nine target locations from neuronal ensembles. Decoding accuracy for ketamine-WM sessions for pre-injection (green), early post-injection (blue), and late post-injection periods (orange) (n=3 sessions) for trial epochs. Data points represent decoding accuracy per session. Chance performance is indicated by dashed grey line. Red center lines indicate median, the bottom and top edges of the box indicate the 25th and 75th percentiles. The whiskers extend to non-outlier data points (approximately within 2.7 std) and the outliers are plotted using ‘+’. *<0.05, **<0.01, ***<0.001.

**Figure 5.**
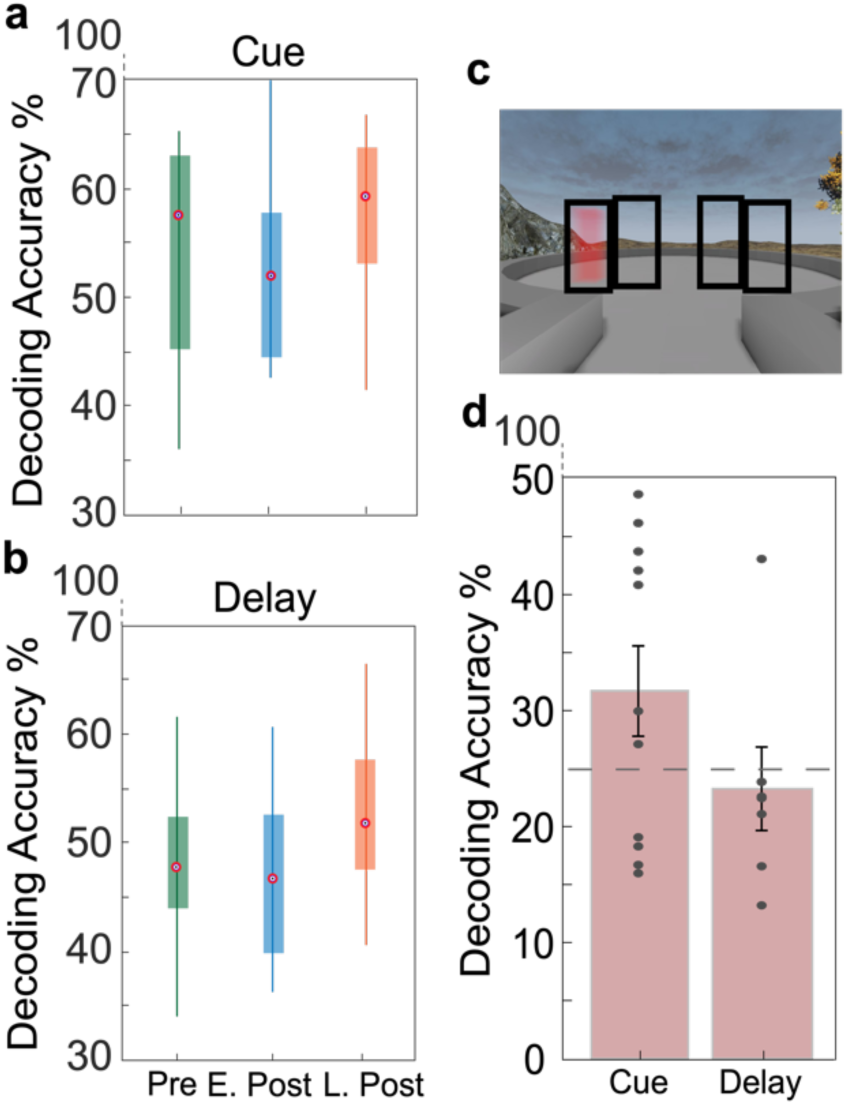
Decoding of eye position and target location during eye fixation. **a**, Comparison of decoding accuracy for target locations using eye fixation position between pre, early, and late post ketamine-injection periods for the cue epoch. **b**, Comparison of decoding accuracy for target locations using eye fixation position between pre, early, and late post ketamine-injection periods for the delay epoch. Red center lines indicate median, the bottom and top edges of the box indicate the 25th and 75th percentiles. The whiskers extend to non-outlier data points (approximately within 2.7 std) and the outliers are plotted using ‘+’. **c**, Four screen regions outlined in black were used in decoding eye position analysis shown in d. **d**, Decoding accuracy for eye position on screen from neuronal ensembles for cue and delay epochs. Data points represent decoding accuracy per session. Dashed grey line indicated chance level. Error bars are SEM. *<0.05, **<0.01, ***<0.001.

**Figure 6.**
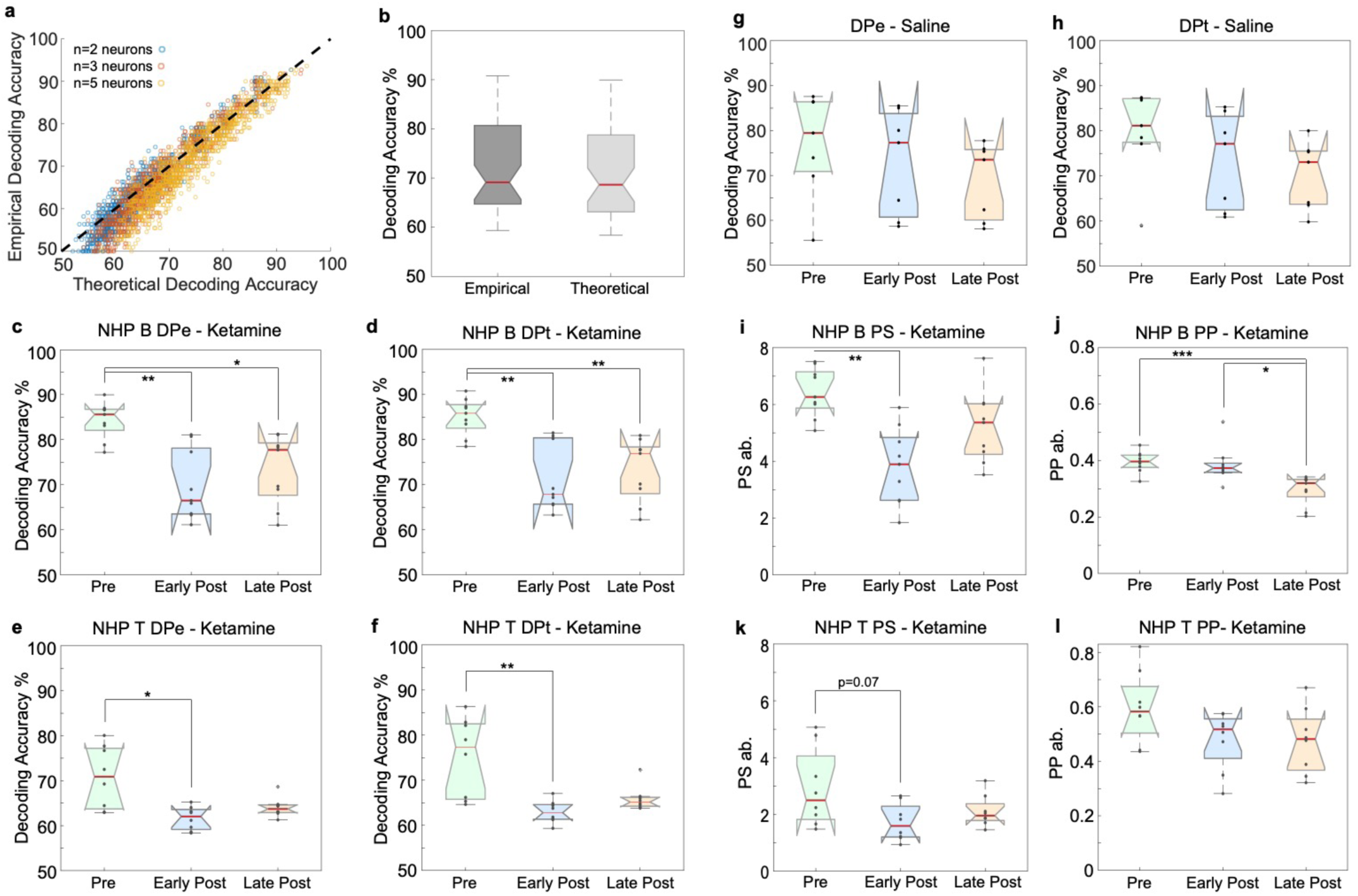
Empirical and theoretical decoding per subject. **a**, Example session showing empirical and theoretical decoding accuracy per neuronal ensemble with decoding above chance level (50%). **b**, Comparison between empirical (DPe) and theoretical (DPt) decoding accuracy for ketamine-WM sessions combined between epochs for the pre-injection period. **c**, DPe for ketamine-WM sessions from NHP B. **d**, DPt for ketamine-WM sessions from NHP B. **e**, DPe for ketamine-WM sessions from NHP T. **f**, DPt for ketamine-WM sessions from NHP T. **g**, DPe for saline-WM sessions. **h**, DPt for saline-WM sessions. **i**, PS for ketamine-WM sessions for NHP B. **j**, PP for ketamine-WM sessions for NHP B. **k**, PS for ketamine-WM sessions for NHP T. **l**, PP for ketamine-WM sessions for NHP T. Data points represent values per session. Red center lines indicate median, the bottom and top edges of the box indicate the 25th and 75th percentiles. The whiskers extend to non-outlier data points (approximately within 2.7 std) and the outliers are plotted using ‘+’. *<0.05, **<0.01, ***<0.001.

**Figure 7.**
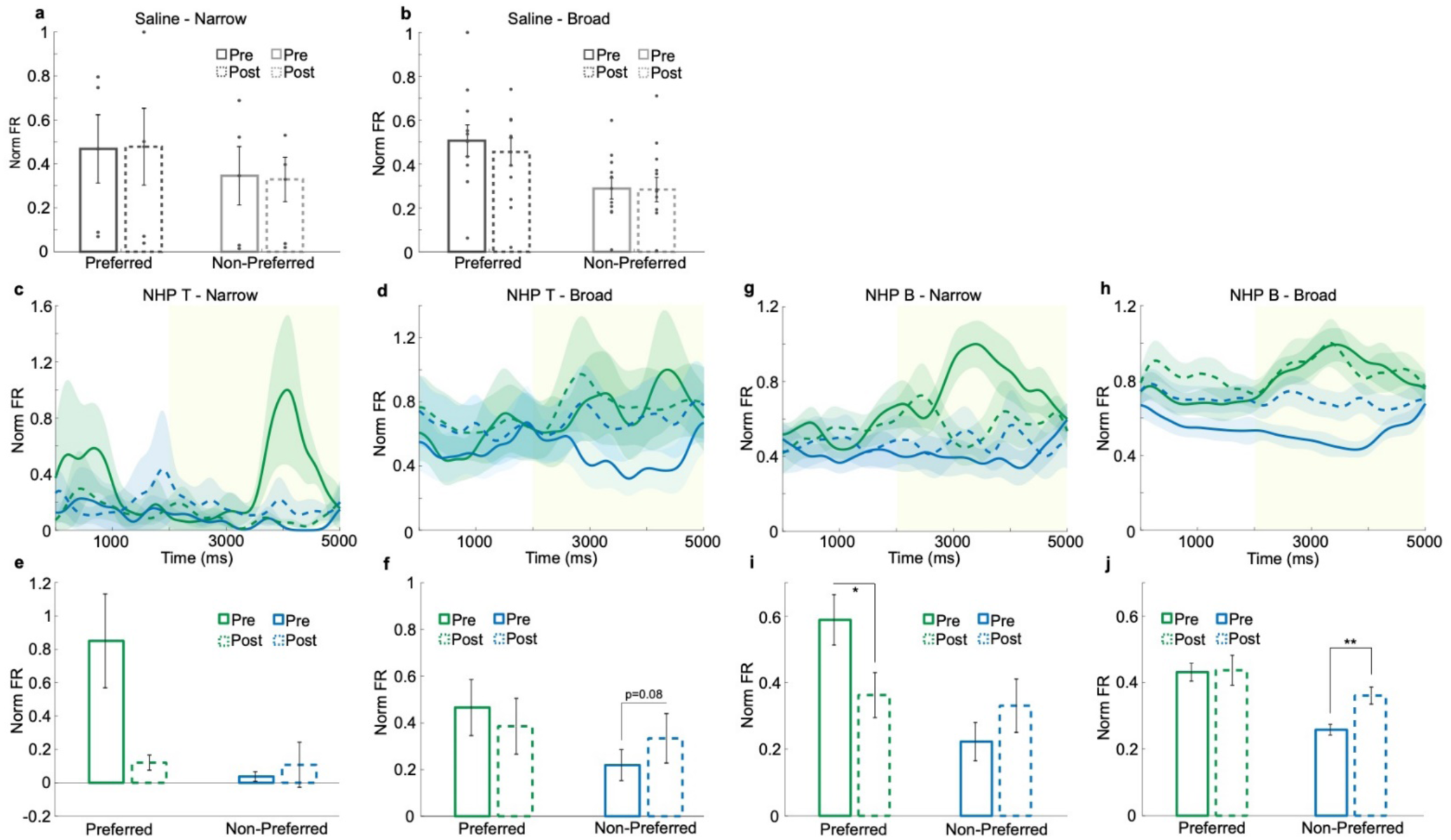
Changes in narrow and broad neuron firing rates per subject. **a**, Firing rates for saline-WM sessions for narrow spiking neurons averaged over the delay epoch for preferred and least-preferred locations. **b**, Firing rates for saline-WM sessions for broad spiking neurons averaged over the delay epoch for preferred locations and least-preferred locations. Data points represent values per electrode array for each session. **c**, Population SDFs for cue and delay (yellow) epochs for delay tuned narrow spiking neurons for NHP T ketamine-WM sessions. **d**, Firing rates for narrow spiking neurons averaged over the delay epoch for preferred and least-preferred locations for pre and early post-injection periods for NHP T ketamine-WM sessions. **e**, Population SDFs for cue and delay (yellow) epochs for delay tuned broad spiking neurons for NHP T ketamine-WM sessions. **f**, Firing rates for broad spiking neurons averaged over the delay epoch for preferred and least-preferred locations for pre and early post-injection periods for NHP T ketamine-WM sessions. **g**, Population SDFs for cue and delay (yellow) epochs for delay tuned narrow spiking neurons for NHP B ketamine-WM sessions. **h**, Firing rates for narrow spiking neurons averaged over the delay epoch for preferred and least-preferred locations for pre and early post-injection periods for NHP B ketamine-WM sessions. **i**, Population SDFs for cue and delay (yellow) epochs for delay tuned broad spiking neurons for NHP B ketamine-WM sessions. **j**, Firing rates for broad spiking neurons averaged over the delay epoch for preferred and least-preferred locations for pre and early post-injection periods for NHP B ketamine-WM sessions. Error bars are SEM. *<0.05, **<0.01, ***<0.001.

**Figure 8.**
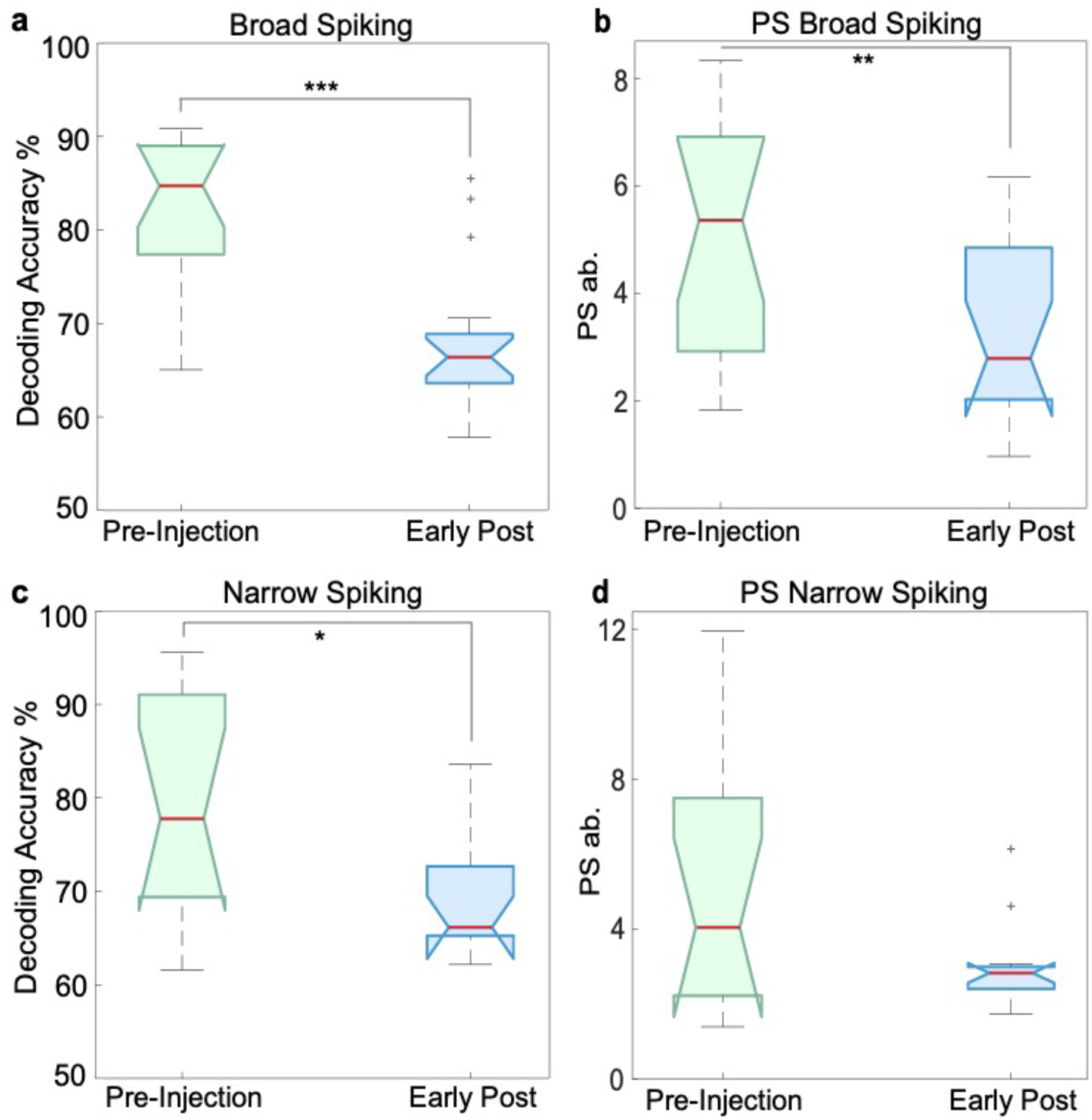
Theoretical decoding and population signal for narrow and broad spiking neurons. **a**, DPt of target location (left/right) from neuronal ensembles of broad spiking neurons. Decoding accuracy over the delay epoch for ketamine-WM sessions (*n*=16) compared between pre and early post-injection periods. **b**, PS for ketamine-WM sessions for broad spiking neuronal ensembles compared between pre and early post-injection periods. **c**, DPt of target location from neuronal ensembles of narrow spiking neurons. Decoding accuracy over the delay epoch for ketamine-WM sessions (*n*=12) compared between pre and early post-injection periods. **d**, PS for ketamine-WM sessions using narrow spiking neurons compared between pre and early post-injection periods. Red center lines indicate median, the bottom and top edges of the box indicate the 25th and 75th percentiles. The whiskers extend to non-outlier data points (approximately within 2.7 std) and the outliers are plotted using ‘+’. *<0.05, **<0.01, ***<0.001.

**Figure 9.**
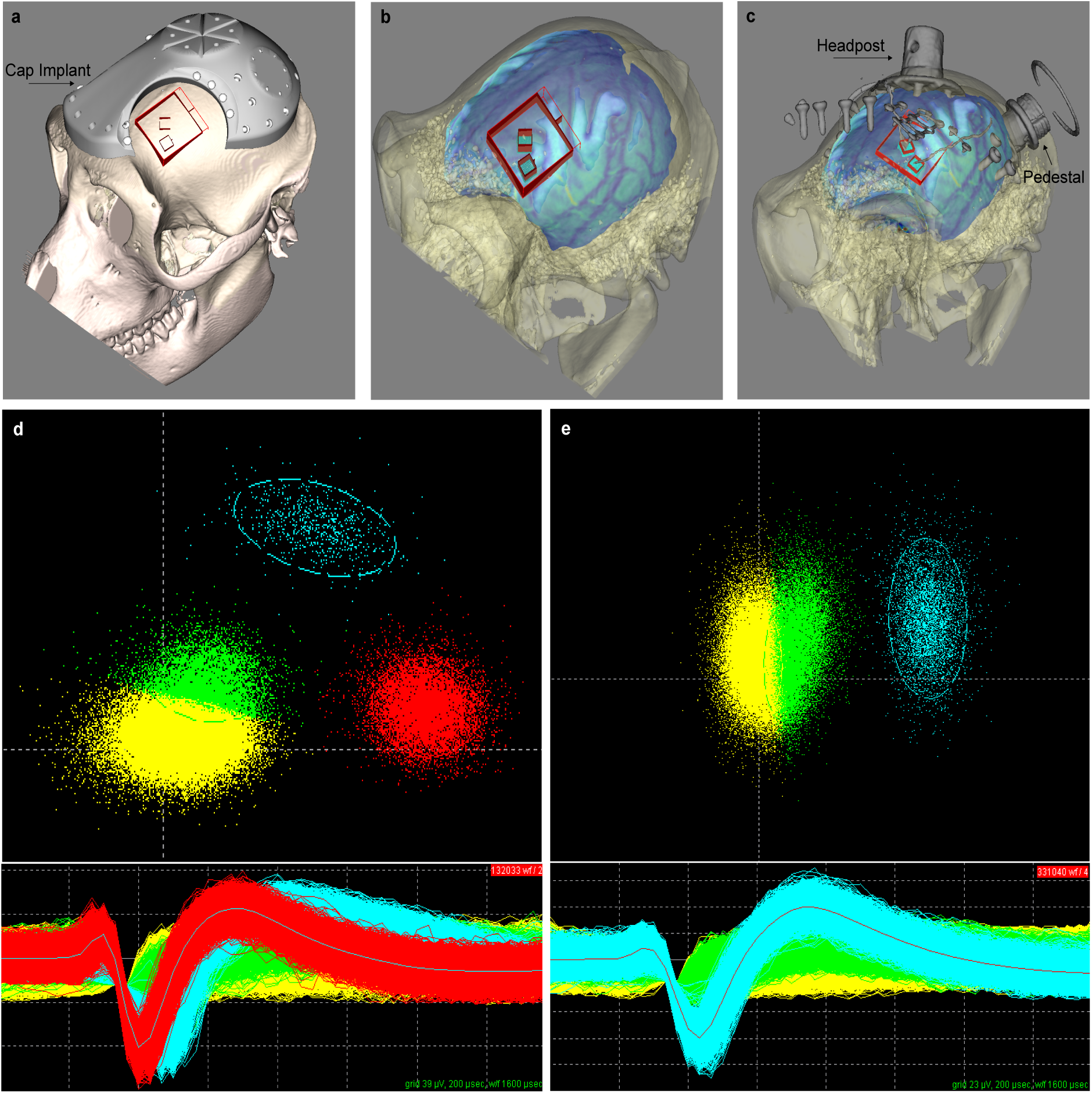
Neural recording and spike sorting. **a**, 3D modeled skull using CT scan from NHP B showing pre-surgical planning conducted in BrainSight. Cap implant depicted as well as planned craniotomy site and planned array implantation site in reference to cap implant. **b**, 3D modeled skull and brain using CT scan and MRI from NHP B showing pre-surgical planning. **c**, 3D modeled skull and brain with overlaid CT imaged cap implant attachments, bone screws, and Utah arrays positioned approximately over the planned array implantation sites. **d**, Spike sorting example from NHP B using Plexon offline sorter. For our analysis, yellow waveforms are considered noise, the green unit would be classified as a multiunit and the blue and red units would be classified as single units. **e**, Spike sorted neuron example from NHP T. The green unit would be classified as a multiunit and the blue unit would be classified as a single unit.

**Figure 10.**
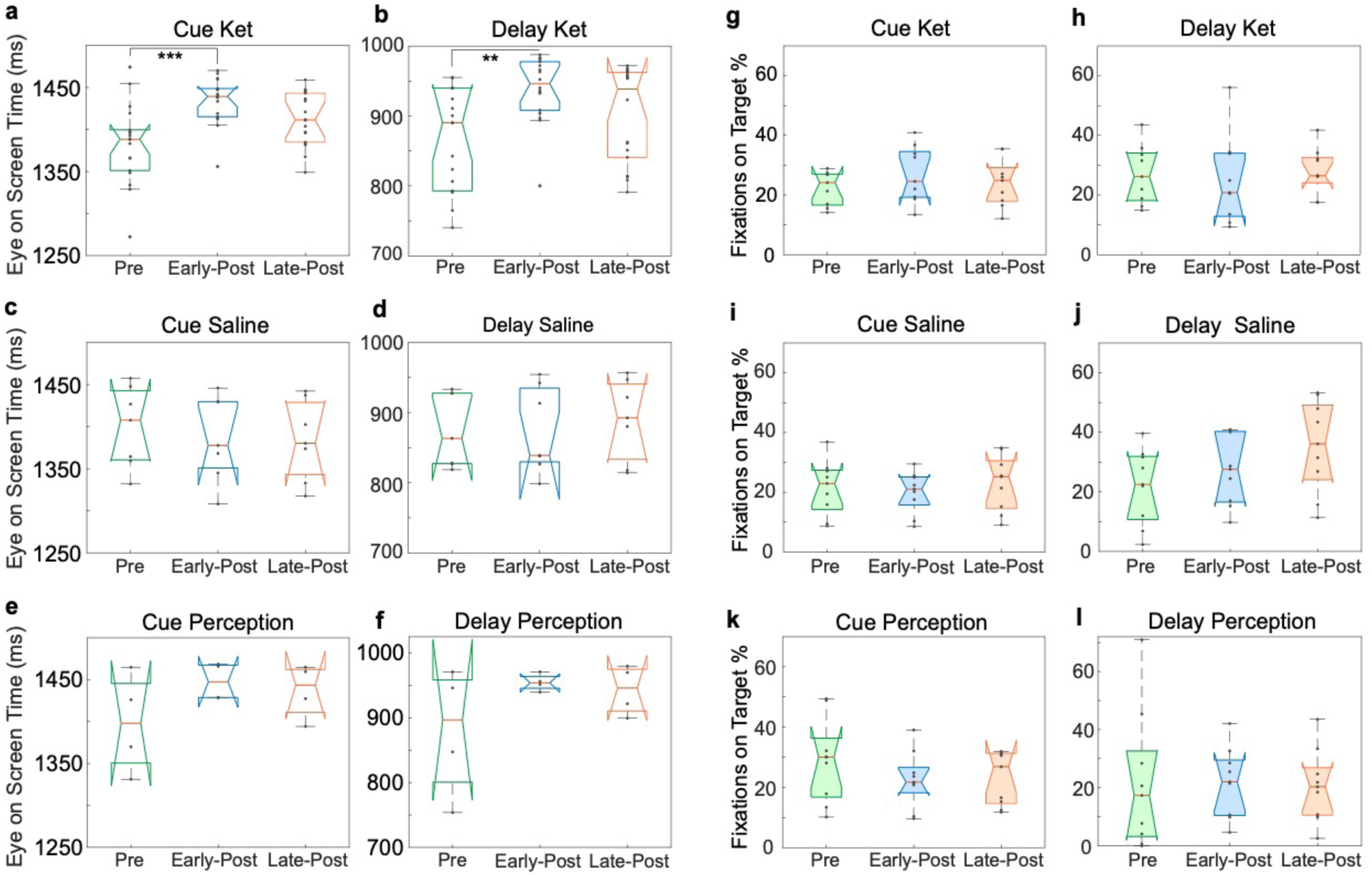
Gaze behaviour. **a**, Duration of eyes on screen during the cue epoch for ketamine-WM sessions. **b**, Duration of eyes on screen during the delay epoch for ketamine-WM sessions. **c**, Duration of eyes on screen during the cue epoch for saline-WM sessions. **d**, Duration of eyes on screen during the delay epoch for saline-WM sessions. **e**, Duration of eyes on screen during the cue epoch for ketamine-perception sessions. **f**, Duration of eyes on screen during the delay epoch for ketamine-perception sessions. Data points represent values per session. **g**, Percentage of fixations on target location during the cue epoch for ketamine-WM sessions. **h**, Percentage of fixations on target location during the delay epoch for ketamine-WM sessions. **i**, Percentage of fixations on target location during the cue epoch for saline-WM sessions. **j**, Percentage of fixations on target location during the delay epoch for saline-WM sessions. **k**, Percentage of fixations on target location during the cue epoch for ketamine-perception sessions. **l**, Percentage of fixations on target location during the delay epoch for ketamine-perception sessions. Data points represent values per target location condition. Red center lines indicate median, the bottom and top edges of the box indicate the 25th and 75th percentiles. The whiskers extend to non-outlier data points (approximately within 2.7 std) and the outliers are plotted using ‘+’. *<0.05, **<0.01, ***<0.001.

## Supplementary Material

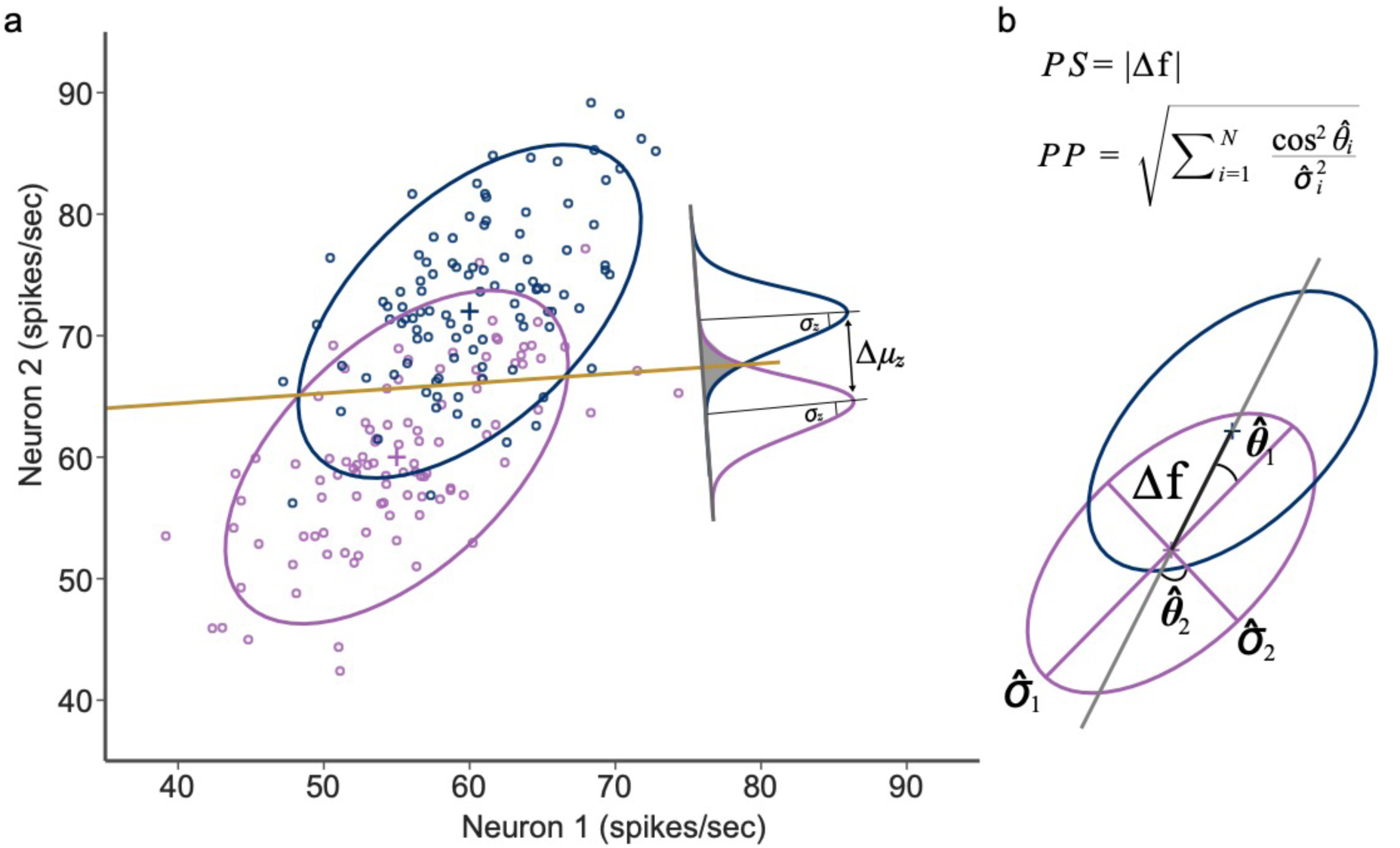
Illustration of population signal and projected precision. **a.** Simulated data illustrates firing rates for two neurons. The covariance matrix and mean activity of the population of neurons determine the shape and location of the ellipsoid for target location 1 (purple) and target location 2 (blue) whereas the yellow line represents a linear classifier that divides the two clouds of data points. **b**. Illustration of population signal and projected precision.

